# The genetic code is very close to a global optimum in a model of its origin taking into account both the partition energy of amino acids and their biosynthetic relationships

**DOI:** 10.1101/2021.08.01.454621

**Authors:** Franco Caldararo, Massimo Di Giulio

## Abstract

We used the Moran’s I index of global spatial autocorrelation with the aim of studying the distribution of the physicochemical or biological properties of amino acids within the genetic code table. First, using this index we are able to identify the amino acid property - among the 530 analyzed - that best correlates with the organization of the genetic code in the set of amino acid permutation codes. Considering, then, a model suggested by the coevolution theory of the genetic code origin - which in addition to the biosynthetic relationships between amino acids took into account also their physicochemical properties - we investigated the level of optimization achieved by these properties either on the entire genetic code table, or only on its columns or only on its rows. Specifically, we estimated the optimization achieved in the restricted set of amino acid permutation codes subject to the constraints derived from the biosynthetic classes of amino acids, in which we identify the most optimized amino acid property among all those present in the database. Unlike what has been claimed in the literature, it would appear that it was not the polarity of amino acids that structured the genetic code, but that it could have been their partition energy instead. In actual fact, it would seem to reach an optimization level of about 96% on the whole table of the genetic code and 98% on its columns. Given that this result has been obtained for amino acid permutation codes subject to biosynthetic constraints, that is to say, for a model of the genetic code consistent with the coevolution theory, we should consider the following conclusions reasonable. (i) The coevolution theory might be corroborated by these observations because the model used referred to the biosynthetic relationships between amino acids, which are suggested by this theory as having been fundamental in structuring the genetic code. (ii) The very high optimization on the columns of the genetic code would not only be compatible but would further corroborate the coevolution theory because this suggests that, as the genetic code was structured along its rows by the biosynthetic relationships of amino acids, on its columns strong selective pressure might have been put in place to minimize, for example, the deleterious effects of translation errors. (iii) The finding that partition energy could be the most optimized property of amino acids in the genetic code would in turn be consistent with one of the main predictions of the coevolution theory. In other words, since the partition energy is reflective of the protein structure and therefore of the enzymatic catalysis, the latter might really have been the main selective pressure that would have promoted the origin of the genetic code. Indeed, we observe that the β-strands show an optimization percentage of 94.45%, so it is possible to hypothesize that they might have become the object of selection during the origin of the genetic code, conditioning the choice of biosynthetic relationships between amino acids. (iv) The finding that the polarity of amino acids is less optimized than their partition energy in the genetic code table might be interpreted against the physicochemical theories of the origin of the genetic code because these would suggest, for example, that a very high optimization of the polarity of amino acids in the code could be an expression of interactions between amino acids and codons or anticodons, which would have promoted their origin. This might now become less sustainable, given the very high optimization that is instead observed in favor of partition energy but not polarity. Finally, (v) the very high optimization of the partition energy of amino acids would seem to make a neutral origin of the ability of the genetic code to buffer, for example, the deleterious effects of translation errors very unlikely. Indeed, an optimization of about 100% would seem that it might not have been achieved by a simple neutral process, but this ability should probably have been generated instead by the intervention of natural selection. In actual fact, we show that the neutral hypothesis of the origin of error minimization has been falsified for the model analyzed here. Therefore, we will discuss our observations within the theories proposed to explain the origin of the organization of the genetic code, reaching the conclusion that the coevolution theory is the most strongly corroborated theory.

## Introduction

### The work plane

The origin of the genetic code is one of the most fascinating problems that can be dealt with because representing the birth of the relationship between the genotype and the phenotype, that is to say, of a unique adaptive moment in the history of life, it allowed the emergence of the biological complexity on our planet. Two factors seem to have occurred in the origin of the organization of the genetic code table: biosynthetic relationships between amino acids and their physicochemical properties (Di Giulio, 1997b, 2005, 2013; Wong, 2005; Koonin and Novozhilov, 2009, 2017). However, there is no agreement on which of these actually played the decisive role. In fact, there are many studies tending to show that biosynthetic relationships between amino acids were extremely important in determining its structuring (Dillon, 1973; Wong, 1975; Taylor and Coates, 1989; Freeland et al., 2000; Di Giulio, 2004, 2008a, 2017a, 2018b; Di Giulio and Amato, 2009). On the contrary, other works favour the conclusion that it was instead the physicochemical properties of amino acids that heavily affected his organization (Woese et al., 1966; Nelsestuen, 1978; Wolfenden et al., 1979; Sjostrom and World, 1985; Di Giulio, 1989a, 1989b; Lacey Jr. et al., 1992; Freeland et al., 2000; Buhrman et al., 2013; Błaz ej et al., 2016, 2018, 2019). Models have also been proposed that take into account both the above factors simultaneously, in an attempt to better understand the origin of the genetic code organization. (Di Giulio, 1991; Freeland et al., 2000; Gilis et al., 2001; Facchiano and Di Giulio, 2018). To achieve this, we first need a method that identifies, as far as possible, the physicochemical or biological properties of amino acids that actually played a key role in organizing the code. Many studies seem to have proved that the genetic code is very conservative with respect to polar requirement (Woese, 1965, 1967; Woese et al., 1966; Di Giulio, 1989a; Haig and Hurst, 1991; Freeland and Hurst, 1998; Butler et al., 2009; Buhrman et al., 2013; Błaz ej 2016, 2018, 2019). Therefore, at present we might observe with Vetsigian et al. (2006) that: “although we do not know what defines amino acid ‘similarity’ in the case of the code, we do know one particular amino acid measure that seems to express it quite remarkably in the coding context. That measure is amino acid polar requirement.” In spite of this, in the first part of this study, we have tried again, but through a different approach, to identify the amino acid properties better related to the structure of the genetic code.

Also regarding the level of optimization actually achieved by physicochemical properties during the origin of the genetic code there is no unanimous consensus (Di Giulio, 2013). In fact, there are works which would indicate that this level is very high (Haig and Hurst, 1991; 1998 Freeland and Hurst; Butler et al. 2009; Buhrman et al. 2013), thus accrediting the physicochemical theories of the origin of the code (Sonneborn, 1965; Woese, 1965, 1967; Lacey Jr. et al., 1992; Fitch and Upper, 1987; Higgs, 2009). While other studies would argue that physicochemical properties of amino acids have reached only a partial level of optimization, on the contrary, supporting theories such as that of the coevolution that attributes to physicochemical properties only a secondary role, identifying instead in biosynthetic relationships between amino acids the crucial factor that led to the structuring of the genetic code (Wong, 1980; Di Giulio, 1989a, 1989b; Di Giulio et al., 1994; Di Giulio and Medugno, 1998, 1999, 2001).

As a result, we have here considered a model of the origin of the genetic code that takes into account both biosynthetic relationships between amino acids and their physicochemical properties (Di Giulio, 1991; Freeland et al., 2000; Gilis et al., 2001; Facchiano and Di Giulio, 2018). In particular, we have determined the level of optimization achieved during the origin of the genetic code, studying the optimization of the physicochemical or biological properties of amino acids not in the set of all possible codes of amino acid permutation (Di Giulio, 1989a) but only in a sub-set subject to the constraints imposed by biosynthetic relationships between amino acids (Facchiano and Di Giulio, 2018). Indeed, operating in this sub-set - with a limited number of elements, but having a very high biological meaning - we hope to be able to make more understandable the crucial steps that led to define the origin of the organization of the genetic code.

In short, using a spatial autocorrelation index and analyzing a large list of amino acid physicochemical and biological properties, we try to: (i) identify the property that best correlates with the organization of the genetic code, without any constraint other than the preservation of the genetic code’s codon block structure; (ii) identify the properties that are best optimized in the restricted set of amino acid permutation codes subject to the biosynthetic constraints, evaluating their optimization separately on columns, rows, and the entire table of genetic code. All this in the hope of being able to: (i) clarify the role actually played by physicochemical properties of amino acids during the origin of the organization of the code; (ii) discriminate between the different theories proposed to explain its origin; and (iii) help to make the crucial steps that led to the structuring of the genetic code more understandable.

### The hypothesis: enzyme catalysis as a determining factor in the evolution of the genetic code

One of the reasons that could have led to triggering the origin of the genetic code might have been that of enzyme catalysis, because the genetic code coding primarily for proteins, essentially catalysts, would seem not to be nothing more than the evolution of coded catalysis (Di Giulio, 2003). In other words, enzyme catalysis - one of the fundamental characteristics of living organisms (Di Giulio, 2015) - may have been the main selective pressure that promoted the origin of the genetic code and determined its structuring (Wong, 1976, 1980; Di Giulio, 1996, 1997a, 2003, 2015). This hypothesis could, among other things, allow to explain the birth of peptidyl-tRNA – the key intermediary of protein synthesis, otherwise not easily selectable (Wong, 1991; Wong and Xue, 2002; Di Giulio, 1997a, 2003, 2007, 2015) – if it were assumed that at least part of catalysis during the origin of the genetic code had to be performed by RNAs covalently linked to peptides, i.e. “peptidyl-tRNAs” (Ageno, 1981; Wong, 1991; Wong and Xue, 2002; Di Giulio, 1997a, 2003, 2007, 2015). This would also allow a first form of coding to evolve naturally, admitting that the interaction between RNA-peptidated (Di Giulio, 1997a, 1997b, 2003, 2007, 2008b, 2015) – which represented a rudimentary form of protein synthesis (Di Giulio, 1997a, 1997b, 2003, 2007, 2008b, 2015) – could be coded by pre-mRNAs, which would in fact have codified sequences of RNA-peptidated interactions (Di Giulio, 2015). From these precursors eventually emerged the actual mRNAs and then the genetic code (Di Giulio, 2015).

It is clear that such a model, while, on the one hand, would be able to elegantly explain the *raison d’être* of the genetic code (Di Giulio, 2008b), on the other hand it would deny any form of stereochemical interaction between codons or anticodons and amino acids as a mechanism for the origin of the code (Di Giulio, 2008a). Indeed, such a model would tie the origin of the genetic code to a great biological complexity that would have facilitated its triggering (Di Giulio, 1996, 1997a, 2003, 2015). Although some authors have seen in this biological complexity an obstacle to the origin of the code (Kun and Radvinyi, 2018), on the contrary, in our view of an evolutionary stage with a relatively high biological complexity would represent a necessary condition for the origin of the genetic code. In other words, we believe that the genetic code cannot originate in a protocell with low or very low biological complexity precisely because, under these conditions, its origin would be completely useless and unnecessary (Di Giulio, 1996, 1997, 2003, 2015). So the catalysis mediated by RNAs bonded covalently to peptides might have been the only propulsive force that would have determined the structuring of the genetic code (Wong, 1976, 1991, 1980; Di Giulio, 1996, 1997a, 2003, 2015).

If this were true, then at a certain evolutionary stage of the origin of the genetic code the physicochemical or biological properties of amino acids should have played an important role in helping the evolution of the primordial enzymatic catalysis. Thus, if this were indeed found for some properties particularly functional to catalysis, this data could provide clarification on the allocations of amino acids within the table of the genetic code. In any case, one would still expect to find the physicochemical properties of amino acids in the table of the genetic code if indeed enzyme catalysis had been the main selective pressure of its origin. A prediction, evidently, shared with the physicochemical theories of the origin of the genetic code (Sonneborn, 1965; Woese, 1965, 1967; Lacey Jr. et al., 1992; Fitch and Upper, 1987; Higgs, 2009) although for completely different reasons (Di Giulio, 1997b). Thus, the physicochemical properties of amino acids reflected in the genetic code (Woese et al., 1966; Alff-Steinberger, 1969; Jungck, 1978; Nelsestuen, 1978; Wolfenden et al., 1979; Sjostrom and World, 1985; Di Giulio, 1989a, 1989b; Lacey Jr. et al., 1992; Freeland et al., 2000; Buhrman et al., 2013; Błaz ej et al., 2016, 2018, 2019; Wnętrzak et al., 2018) might be an expression of enzyme catalysis and may not reflect physicochemical constraints imposed by its origin. So, it is also to try to give some answer to this problem that we have undertaken this study.

To recap, since we are convinced of the substantial correctness of the coevolution theory of the origin of the genetic code (Wong, 1975; Di Giulio, 2008a) - which assumes, among other things, that enzyme catalysis was precisely the main selective pressure that promoted the origin of the code (Wong, 1976, 1980; Di Giulio, 1996, 1997a, 2003, 2015) - we tried to understand if the physicochemical properties of amino acids observed in the genetic code are reflective of enzyme catalysis or whether they may otherwise be an expression of interactions of codons or anticodons and amino acids or other types of interactions or physicochemical mechanisms that would have promoted the origin of the genetic code.

## Materials and Methods

### The database

Here we refer to the list of 544 physicochemical and biological properties for the 20 amino acids imported in the R package ‘seqinr’ (version 4.2-5; 2020-12-17; http://seqinr.r-forge.r-project.org/) from release 9.1 (Aug 2006) of the aaindex1 database (Kawashima and Kanehisa, 2000). After deleting binary variables and properties containing NA values, we extended the database to include the Original Polar Requirements (Woese et al., 1966), Updated Polar Requirements (Mathew and Luthey-Shulten, 2008), Mobilities of amino acids on chromatography paper (Aboderin, 1971) and the Effective Mean Energy (Miyazawa-Jernigan, 1996), so that a total of 530 properties for the 20 amino acids were analyzed.

### Computational methods

Results presented here were obtained using R programming language and free software environment (version 4.0.3, https://cran.r-project.org/) for statistical computing (R Core Team, 2020), using - in addition to the ‘seqinr’ package (Charif and Lobry, 2007) described above - the spdep library (version 1.1-5), a collection of tests for spatial autocorrelation (Bivand et al., 2018), and the ‘gtools’ package (version 3.8.2), various R programming tools including permutations and combinations (Warnes et al., 2020). R codes written by the authors can be obtained upon request.

### The spatial autocorrelation index

To find out which database properties are best correlated with the organization of the genetic code, we use an autocorrelational approach. In actual fact, if you keep the blocks of synonymous codons unchanged, which in the genetic code encode for the 20 amino acids and for the termination signal, you can locate the neighbours for each block and assign weights that measure the intensity of the relationship between pairs of blocks. In other words, blocks of synonymous codons (B_1_, B_2_, …, B_20_) coding for amino acids are seen as locations or spatial units, so - from an operational point of view - the notion of spatial proximity is formally expressed through a 20 x 20 weight matrix W, whose generic off-diagonal element, wij, represents the relationship between B_i_ and B_j_. In particular, in our model, w_ij_ represents the number of pairs that differ exactly by one base, among all those that can be formed by choosing the first codon in B_i_ and the second in B_j_. It should also be noted that the blocking of termination codons is not taken into account (however, it indirectly manifests its action in the smaller number of neighbors assigned – on the same terms – to the locations adjacent to it). Of course, depending on how the 20 amino acids are uniquely assigned to the blocks, the corresponding values (x_1_, x_2_, …, x_20_) of a given physicochemical or biological property can take up different places in space. In this context, spatial autocorrelation refers to the relationship between spatial proximity between blocks of synonymous codons and the numerical similarity between values of amino acids coded by them. Therefore, spatial autocorrelation indices allow us to measure the correlation of a specific physicochemical or biological property with itself, taking into account the spatial arrangement of its values assigned to the synonymous codons blocks.

Our measurement of spatial autocorrelation is based on the Moran I statistic (Moran, 1948, 1950; Cliff and Ord, 1973, 1981), arguably the most well known and most widely applied statistic for testing spatial autocorrelation. We denote by the observed value of a given physicochemical or biological property in the block B_i_ (i=1,2, …, n), by the mean of all the observed 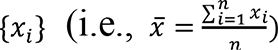 and by w_ij_ the generic element of the spatial weight matrix W. Using this notation, the Moran I coefficient of global spatial autocorrelation can be defined as:

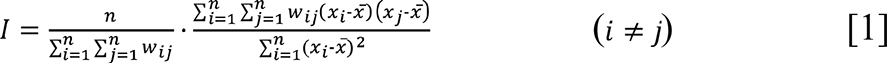

Thus, we see that the numerator is a measure of covariance among the {*x_i_*} and the denominator is a measure of variance. It should also be noted that this coefficient is based upon the cross-products of the deviations from the mean and that a key role is played by the weighting matrix W. Values of the index usually fall in the [-1, 1] interval and can be interpreted by analogy with Pearson correlation coefficient. In our context, Moran I is positive when nearby blocks of synonymous codons tend to encode similar amino acids, negative when the coded amino acids tend to be more dissimilar than one might expect, and approximately zero when they are arranged randomly and independently in their relative blocks.

Under fairly wide assumptions (Cliff and Ord, 1971), the sampling distribution of Moran’s I statistic is known to be asymptotically normal. In such a case, the expected value and the variance of *I* can algebraically be evaluated (Cliff and Ord, 1981), under the null hypothesis of spatial randomness, either by: (i) assuming that the observed {*x_i_*} (i=1,2,…n) are random independent drawings from a normal population (assumption N - normality); and (ii) considering the observed value of *I* with reference to the set of all the possible values which *I* could take on if the {*x_i_*} values were repeatedly randomly permuted around the n spatial units (assumption R - randomisation).

When normality cannot be assumed, a reference distribution under the null hypothesis of spatial randomness is empirically generated for *I* by a Monte Carlo simulation that generates a large number of permutations of the observed {x_i_} over all locations and by re-computing Moran I for each sample. In our case, let I_code_ be the Moran I obtained for the (not randomized) observed {x_i_} in the standard genetic code, if we denote by N_sim_ the number of permutations in the simulation and by N_GE_ the number of times I under simulation was greater than or equal to I_code_, than a pseudo-significance can be computed as:

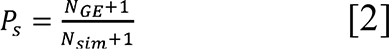

In the event that the set of permutations permutations is not too large, then - through a complete enumeration procedure - we can evaluate the value of index *I* for each element of the collection and identify as well the element with the maximum value of the index.

Taking into account all possible single base and sense changes between the codons of the genetic code, we indicated with *I* the global Moran index of spatial autocorrelation associated with the spatial matrix of W weights, as defined above. But, by specializing the analysis to sense and single base changes involving only the columns of the code, or only those of the rows, or those involving changes only in the first, or only in the second position of codons, we come in a similar way to the definition of other global spatial autocorrelation indices (I_C_, I_R_, I_1_ and I_2_) associated with different weight matrices (W_C_, W_R_, W_1_, W_2_), respectively.

Lastly, the spatial weight matrix is often row-standardized (each weight is divided by its row sum), but “there is no mathematical or statistical requirement for this” (Anselin, 1988, page 23). Undoubtedly, this strategy facilitates the interpretation of the statistics: for instance, it effectively enables us to interpret the spatial lag term as a spatially-weighted average of neighbouring values. Mainly, row-standardization is used to mitigate the effect due to the unequal number of neighbours. But, despite its popularity, row-standardization has disadvantages. Primarily, it upsets the internal weighting structure of the spatial weight matrix, realistically destroying any symmetry and, perhaps, risking to distort the very nature of the spatial interactions we are modelling. Moreover, due to row-standardization each row sums 1 and the total influence on each spatial unit becomes the same. All this can hardly be considered neutral. In our model, in practice, row-standardization penalizes blocks with a high (>2) number of synonym codons, favoring the remaining, especially when these are contiguous to the termination codons and/or included in the same codon box. Thus, due to the effect of row-standardization alone, blocks with a small number of synonymous codons will have a stronger impact on the global Moran I. On the contrary, the natural structure of the synonymous codon block system requires that in our model the size of the blocks must be taken into account adequately, since it reflects an essential trait, important for the evolution of the genetic code: for example, a significant correlation between the number of codons and the molecular weight of amino acids is known (Hasegawa and Miyata, 1980; Taylor and Coates, 1989; Di Giulio, 1989b, 1997b, 2005; Dufton, 1997). Therefore, we believe that results obtained with non-standardized matrices are to be considered more reliable from a strictly evolutionary point of view. On the other hand, row-standardization leads to a substitution matrix (whose entries represent transition probabilities between synonymous codon blocks) which certainly has its biological relevance as well, since it highlights the mutational effects between blocks that encode for different amino acids. For these reasons we present both results, with and without row-standardization.

### Amino acid permutation codes and genetic code optimality

In this work, we will look at two models. In the first, for each specific amino acid property, we were going to consider – as a set of admissible permutations - all 20! possible permutations of its values (Di Giulio, 1989a). In the second model, on the other hand, we will operate in the restricted set of amino acid permutations subject to biosynthetic constraints, in which the amino acids of the same biosynthetic family are allowed to permutate exclusively on the blocks of synonymous codons really assigned by the genetic code to the family itself (Facchiano and Di Giulio, 2018). The biosynthetic constraints considered are derived from the following biosynthetic classes (Wong, 1975; Taylor and Coates, 1989; Di Giulio, 2008a, 2018; Facchiano and Di Giulio, 2018):

1. Ser, Gly, Cys, Trp;
2. Phe, Tyr;
3. Val, Leu, Ala;
4. Glu, Gln, His, Arg, Pro;
5. Asp, Asn, Lys, Thr, Met, Ile.

Since the number of amino acid permutations (Di Giulio, 1989a) allowed by this model is 4!2!3!5!6!= 24,883,200 (see also, Facchiano and Di Giulio, 2018), it is now possible to evaluate index *I* for each element of the set. In particular, you can know the number (Nsg) of all values of *I* strictly greater than I_code_ and identify the maximum value (I_max_). As a result, an estimate of the level of optimization achieved by the genetic code can be made using the optimization percentage (Opt) calculated - in analogy with the percentage of minimization (Wong, 1980; Di Giulio, 1989b) - by the expression:

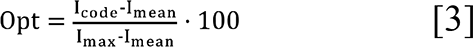

where I_code_ is the Moran’s I value of the genetic code, I_mean_ is the average value of the index, and I_max_ is its maximum value.

All the R codes written by the authors can be obtained upon request.

## Results

### Identification of physicochemical properties of amino acids that best correlate with the organization of the genetic code

We performed calculations of Moran’s I statistics (Moran, 1948, 1950; Cliff and Ord, 1973, 1981) of global spatial autocorrelation for all 530 physicochemical and biological properties for the 20 amino acids of the database described above, with the aim of identifying properties which had a more significant spatial autocorrelation. In this sense, we can briefly refer to these properties as the best correlated with the structure of the genetic code. Particularly, in a first scan, for each amino acid property we analytically computed five global Moran’s statistics (I, I_1_, I_2_, I_c_, and I_r_), using both row-standardized and non-row-standardized spatial weight matrices. Namely, this has been made or considering all possible changes of single base and of sense between codons of the genetic code (I), or those involving only the columns (I_c_), or only those of the rows (I_r_), or those involving changes only in the first (I_1_), or only in the second (I_2_) position of codons (see Materials and Methods). In this way, it is possible to obtain a first overview of the behavior of the various indices, by observing how these descriptive measures vary from one variable to another. Then, the 50 best-performing properties with respect to global I index (corresponding to the largest zeta values, analytically estimated under the randomization assumption) were chosen for a more careful assessment of the significance of observed values of the various indices by simulation experiments that randomize the values of a given property over the synonymous codon blocks. Thus, for any given spatial weighting scheme, a permutation test was performed to establish the rank (and the pseudo p-value) of the observed statistic of the genetic code in relation to the simulated values, by using 10^7^ (= 9999999 + 1) random permutations, sampled from all 20! possible (Di Giulio, 1989a), for the more significant indices (I, I_1_ and I_c_) and 10^5^ (=99999 + 1) random permutations for the remaining indices (I_2_ and I_r_). Finally, properties were then ordered according to the non decreasing pseudo p-values relative to the global I index. Results of the top 20 properties are presented in table 1 (without row-standardizing) and table 2 (with row-standardized spatial weight matrices).

**Table 1.**
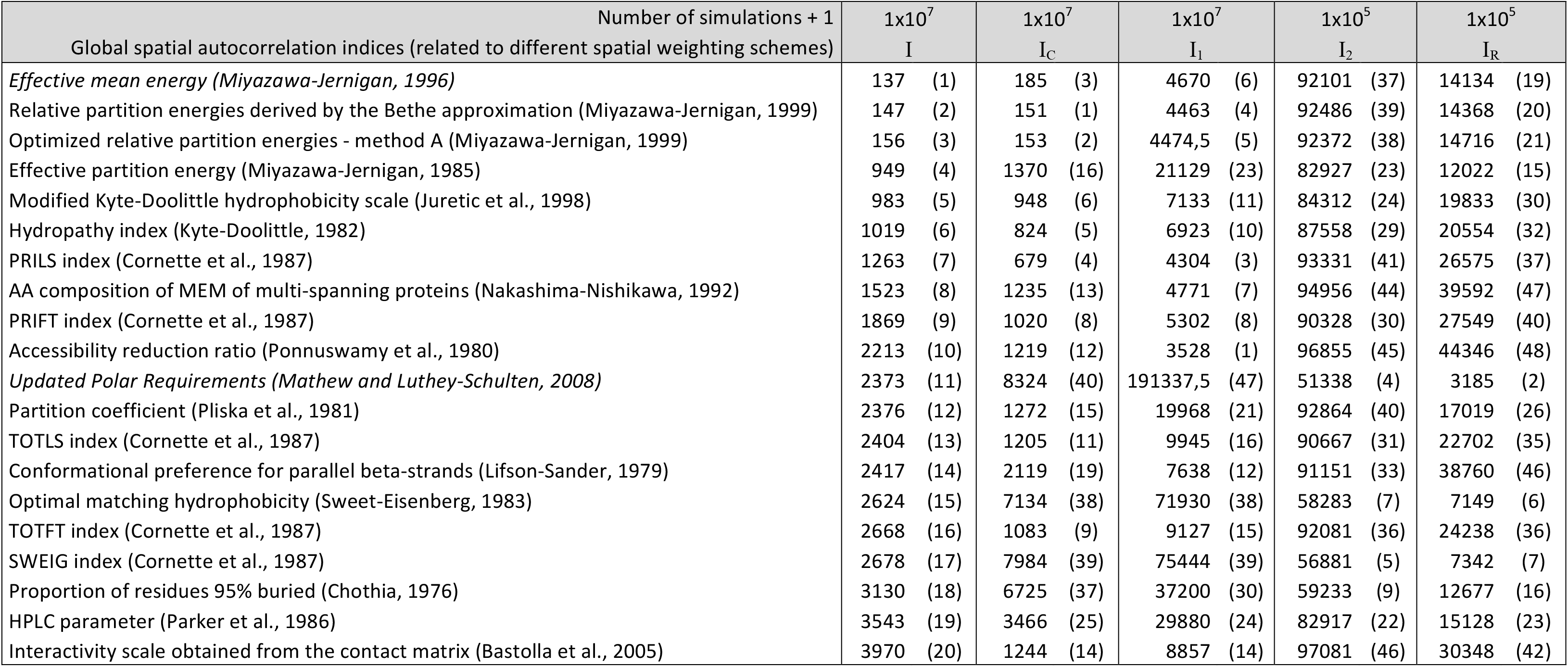
The 20 most significantly auto-correlated amino acid properties from the AAindex database (Kawashima and Kanehisa, 2000). Results of Moran’s I Monte Carlo tests. Each simulation generates a large number of permutations (over all synonymous codons blocks) of the values observed in the genetic code for a given amino acid property and recalculates Moran’s I for each sample. The pseudo p-value of the test is reported, multiplied by the number of simulations +1, for each spatial weighting scheme. In other words, the numbers indicate how many permutation codes were found in the simulation that have the corresponding Moran’s I equal to or greater than the genetic code (also included in the count). The numbers enclosed in parentheses refer to the rank among the 50 analyzed properties. All spatial weight matrices here are not row-standardized. The italic writing emphasizes that the property was added for comparison to the AAindex database.

**Table 2.** The 20 most significantly auto-correlated amino acid properties from the AAindex database (Kawashima and Kanehisa, 2000). Results of Monte Carlo tests of Moran’s I computed using row-standardized spatial weight matrices. Each simulation generates a large number of permutations (over all synonymous codons blocks) of the values observed in the genetic code for a given amino acid property and recalculates Moran’s I for each sample. The pseudo p-value of the test is reported, multiplied by the number of simulations +1, for each spatial weighting scheme. In other words, the numbers indicate how many permutation codes were found in the simulation that have the corresponding Moran’s I equal to or greater than the genetic code (also included in the count). The numbers enclosed in parentheses refer to the rank among the 50 analyzed properties. The italic writing emphasizes that the property was added for comparison to the AAindex database.

| Number of simulations + 1<br>Global spatial autocorrelation indices (related to different spatial weighting schemes) | 1x10<sup>7</sup><br>I | 1x10<sup>7</sup><br>I<sub>C</sub> | 1x10<sup>7</sup><br>I<sub>1</sub> | 1x10<sup>5</sup><br>I<sub>2</sub> | 1x10<sup>5</sup><br>I<sub>R</sub> |
| --- | --- | --- | --- | --- | --- |
| Optimized relative partition energies - method A (Miyazawa-Jernigan, 1999) | 31 (1) | 257 (3) | 2182 (8) | 75545 (27) | 8011 (23) |
| *Effective mean energy (Miyazawa-Jernigan, 1996)* | 41 (2) | 269 (4) | 2290 (9) | 75559 (28) | 7710 (21) |
| Relative partition energies derived by the Bethe approximation (Miyazawa-Jernigan, 1999) | 50 (3) | 208 (1) | 2122 (7) | 75806 (29) | 7735 (22) |
| *Updated Polar Requirements (Mathew and Luthey-Schulten, 2008)* | 62 (4) | 692 (11) | 46711 (40) | 46363 (9) | 1548 (1) |
| Modified Kyte-Doolittle hydrophobicity scale (Juretic et al., 1998) | 166 (5) | 255 (2) | 747 (2) | 84085 (37) | 14045 (35) |
| Hydropathy index (Kyte-Doolittle, 1982) | 248 (6) | 271 (5) | 573 (1) | 90582 (42) | 16029 (40) |
| Effective partition energy (Miyazawa-Jernigan, 1985) | 250 (7) | 761 (12) | 8505 (22) | 78624 (31) | 6971 (18) |
| Polarity (Grantham, 1974) | 335 (8) | 2299 (24) | 47139 (41) | 56949 (13) | 2744 (5) |
| Average side chain orientation angle (Meirovitch et al., 1980) | 342 (9) | 621 (9) | 2773 (11) | 73558 (26) | 8065 (24) |
| Optimized relative partition energies - method D (Miyazawa-Jernigan, 1999) | 432 (10) | 3374 (30) | 25148 (30) | 50313 (12) | 4078 (10) |
| PRILS index (Cornette et al., 1987) | 563 (11) | 559 (8) | 1483 (4) | 85795 (39) | 17663 (43) |
| PRIFT index (Cornette et al., 1987) | 576 (12) | 689 (10) | 1250 (3) | 79303 (32) | 17015 (41) |
| Polar requirement (Woese, 1973) | 689 (13) | 3198 (28) | 119226 (46) | 57110 (14) | 1918 (3) |
| Optimized relative partition energies - method B (Miyazawa-Jernigan, 1999) | 797 (14) | 2935 (27) | 19468 (26) | 57674 (15) | 6145 (16) |
| *Original Polar Requirements (Woese et al., 1966)* | 842 (15) | 5542 (39) | 170948 (48) | 48914 (11) | 1613 (2) |
| Optimal matching hydrophobicity (Sweet-Eisenberg, 1983) | 1052 (16) | 8888 (44) | 59732 (42) | 40601 (7) | 3927 (9) |
| Proportion of residues 95% buried (Chothia, 1976) | 1116 (17) | 2456 (25) | 6185 (17) | 70825 (24) | 12322 (32) |
| Hydropathy scale based on self-information values in the two-state model (9% accessibility) (Naderi-Manesh et al., 2001) | 1161 (18) | 7493 (43) | 26857 (32) | 42870 (8) | 4297 (11) |
| SWEIG index (Cornette et al., 1987) | 1184 (19) | 9743 (45) | 65911 (43) | 39250 (6) | 3875 (8) |
| Direction of hydrophobic moment (Eisenberg-McLachlan, 1986) | 1197 (20) | 302 (6) | 3489 (13) | 99227 (50) | 15923 (38) |

First, taking into account index I, it must be said that all properties listed in tables 1 and 2 have - as widely expected - pseudo p-values so small that it is completely unlikely that the data can support the null hypothesis of spatial randomness. That said, we can consider the pseudo p-values as “exploratory tools”, capable of providing us with indications if a given hypothesis is able to provide an adequate description of the data or if new ones and more plausible alternatives should be investigated. In particular, at the various amino acid properties analyzed, we can consider the pseudo p-values as a computational summary of the extremeness of the observed statistic for the genetic code with reference to the distribution of the simulated random permutation codes.

The data reported so far in the literature support the hypothesis that polarity is the property best correlated with the structure of the code (Woese, 1965, 1967; Woese et al., 1966; Di Giulio, 1989a, 1989b; Haig and Hurst, 1991; Freeland and Hurst, 1998; Butler et al., 2009; ej et al., 2016, 2018, 2019). On the other hand, by analyzing both Tabs. 1 and 2, the observation that is evident is that it is rather the measures of the partition energy of amino acids (Miyazawa-Jernigan, 1985, 1996, 1999) that achieve the best performance, that is to say, to seize the ranks smaller ones. Particularly in Tab. 1 (where, in our opinion, the most reliable situation is represented from a strictly biological point of view, having been used non-standardized spatial weight matrices) appears only in 11^th^ place a single measure of polarity: Updated Polar Requirements (Mathew and Luthey-Schulten, 2008). Now, while it is true that in Tab. 2 (standardized weight matrices) there are four measures of polarity, on the other hand, it should be noted that in both tables, three measures of the partition energy of amino acids are the first to win the top ranks: the relative partition energies derived by the Bethe approximation (Miyazawa-Jernigan, 1999), effective mean energy (Miyazawa-Jernigan, 1996) and optimized relative partition energies - method A (Miyazawa-Jernigan, 1999). In conclusion, it would seem that the partition energy of amino acids is the property best correlated with the structure of the genetic code and not the polarity of amino acids. Therefore, although conclusions of significance tests must always be confirmed by further measurements, at present it seems that our findings question the adequacy of the polarity model, suggesting a more plausible one based on the partition energies. An even worse behavior than that of polarity is observed for the hydration potential of amino acids (Wolfenden et al., 1979) which does not appear to be present in Tabs. 1 and 2, and not even within the first fifty ranks, although some analyses (Wolfenden et al., 1979) indicated that it was strongly correlated with the organization of the genetic code.

When analyzing the Tabs. 1 and 2, while it is appropriate to compare the values of a given index varying from one variable to another, it is incorrect to directly compare the results of two different indices, since they correspond to different spatial weighting schemes. Nevertheless, it is clear that the general trend is that most of the optimization of the genetic code is relegated to the columns (Ic) and this would be consistent with the observations reported in the literature that it is mainly the columns of the genetic code that are well-accommodated by physicochemical properties of amino acids (Nelsestuen, 1978; Wolfenden et al., 1979; Sjostrom and World, 1985; Di Giulio, 1989b, 2017a, 2018b; Chiusano et al., 2000).

The contrast between the results of the two tables is not very marked, since their only difference consists in the row-standardization of the weighting matrix. They share 12 properties which seem roughly to show the same trend, since the correlation coefficient between their ranks is highly significant (r = + 0.900, p=8.3×10^-5^, n = 12). The introduction of the row-standardization of the weighting matrix does not penalize the measures of partition energy at all (on the contrary, in Tab. 2 there are two more, in addition to the 4 shared with Tab. 1) and seems in some way to favor polarity (with the inclusion in the top 20 of 3 other new measures, in addition to the shared one that, moreover, wins a lower rank). The ability of partition energy to be, unlike polarity, relatively insensitive to changes in model assumptions induced by row-standardization, could be interpreted as a further indication of its most reliable suitability to provide an adequate description of the data. Indeed, the partition energy is not only capable of providing the most significant values, but is also able to manifest them both in presence and in the absence of standardization. This suggests that it might have been the partition energy that played the main role in structuring the genetic code, not the polarity. The latter, in actual fact, being penalized by the absence of standardization, is as if it were not able to guarantee a good performance just when considering, without further modification, the natural structure of the synonymous codon blocks, an essential attribute for the evolution of the genetic code (Di Giulio, 1989b, 2005a).

Can we be reasonably certain that we have identified in the partition energy of amino acids the property that best correlates with the organization of the genetic code? We believe that the answer to this question can only be affirmative, of course within the limits imposed by the analysis itself. So, in the next section we will investigate whether this performance of partition energy is also preserved in the codes of amino acid permutations but subject to biosynthetic constraints between amino acids, that is to say, in the set of codes expected from the coevolution theory of the genetic code (Di Giulio, 2016b, 2017a, 2018b).

### Optimality of the genetic code in the coevolution theory framework: partition energies of amino acids and their conformational preference for parallel beta-strands are the only properties to be highly optimized

In the previous section we considered, as the set of allowed permutations for each specific amino acid property, all 20! possible permutations of its values (Di Giulio, 1989a). When, on the other hand, we operate in the set of amino acid permutations subject to biosynthetic constraints, the number of allowed permutations is restricted to only 24883200, thus making it possible to evaluate the Moran’s I index for each element of the set. Thus we are able to compute the optimization percentage (see Materials and methods) reached by the genetic code and the number of codes that have better performance than it. Specifically, by sampling in the subset of restricted permutations, we first estimated the optimization level achieved by the genetic code with respect to the global Moran’s I index of spatial autocorrelation and with reference to each of the 530 properties of the database. Out of the 50 properties with the highest estimates, the I index of global spatial autocorrelation was then exhaustively calculated for each of the 24,883,200 amino acid permutation codes subject to biosynthetic constraints. Finally, only the top 20 properties were selected and the calculations of the remaining indices (I_1_, I_2_, I_C_, I_R_) were performed. The results obtained are shown in Tab. 3 (without row-standardizing) and Tab. 4 (with row-standardized spatial weight matrices).

**Table 3.** The 20 properties with the highest optimization percentages from the AAindex database (Kawashima and Kanehisa, 2000). Optimality of the standard genetic code in the coevolution theory framework. For each global index of spatial autocorrelation, two different optimality measures of the genetic code are reported in the restricted set of permutation codes subject to biosynthetic constraints. The first, Opt, represents the optimization percentage (see Materials and methods) and the second, Nsg, indicates the number of codes that have the relative index value strictly greater than that of the genetic code. The listed properties have the highest optimization percentages, in relation to index I, among all those of the database. All spatial weight matrices here are not row-standardized. The italic writing emphasizes that the property was added for comparison to the AAindex database.

| Global spatial autocorrelation indices<br>(related to different spatial weighting schemes) | I |  | I<sub>C</sub> |  | I<sub>1</sub> |  | I<sub>2</sub> |  | I<sub>R</sub> |  |
| --- | --- | --- | --- | --- | --- | --- | --- | --- | --- | --- |
| Optimality measures | Opt | Nsg | Opt | Nsg | Opt | Nsg | Opt | Nsg | Opt | Nsg |
| Relative partition energies derived by the Bethe approximation (Miyazawa-Jernigan, 1999) | 96,45 | 4429 | 97,74 | 1055 | 92,31 | 10272 | -29,98 | 23271528 | 14,66 | 4868524 |
| *Effective mean energy* (Miyazawa-Jernigan, 1996) | 96,30 | 4555 | 97,65 | 1122 | 91,96 | 10652 | -30,08 | 23294944 | 14,95 | 4781120 |
| Optimized relative partition energies - method A (Miyazawa-Jernigan, 1999) | 96,18 | 4774 | 97,61 | 1165 | 92,09 | 10388 | -29,92 | 23293216 | 14,35 | 4957528 |
| Effective partition energy (Miyazawa-Jernigan, 1985) | 95,94 | 12754 | 97,10 | 3459 | 90,18 | 26836 | -28,11 | 22726704 | 23,98 | 2853940 |
| Conformational preference for parallel beta-strands (Lifson-Sander, 1979) | 95,45 | 13672 | 85,47 | 92526 | 87,79 | 103084 | -19,55 | 17582976 | 9,12 | 9174808 |
| HPLC parameter (Parker et al., 1986) | 92,71 | 2324 | 93,47 | 2264 | 83,33 | 23060 | -21,40 | 22354712 | 15,76 | 3944324 |
| Buriability (Zhou-Zhou, 2004) | 92,26 | 8586 | 94,33 | 4120 | 83,93 | 29604 | -24,57 | 21376424 | 18,70 | 4368108 |
| van der Waals parameter R0 (Levitt, 1976) | 92,22 | 9216 | 73,81 | 95040 | 72,50 | 319968 | 13,97 | 6402240 | 37,83 | 2063232 |
| Interactivity scale obtained by maximizing the mean of correlation coefficient over single-domain globular proteins (Bastolla et al., 2005) | 92,13 | 6032 | 92,77 | 5307 | 87,72 | 17752 | -23,25 | 21302856 | 9,12 | 7510632 |
| Accessibility reduction ratio (Ponnuswamy et al., 1980) | 91,78 | 24944 | 91,21 | 37682 | 92,22 | 15772 | -21,88 | 22819376 | 2,71 | 9420104 |
| Hydropathy index (Kyte-Doolittle, 1982) | 89,27 | 23260 | 91,73 | 10644 | 89,39 | 13840 | -29,60 | 23595216 | 8,89 | 7378624 |
| PRILS index (Cornette et al., 1987) | 89,02 | 38149 | 91,39 | 21751 | 86,97 | 39712 | -22,52 | 22270760 | 4,83 | 8198012 |
| Average surrounding hydrophobicity (Manavalan-Ponnuswamy, 1978) | 88,99 | 45243 | 92,64 | 13996 | 90,10 | 15708 | -32,66 | 24308824 | -2,12 | 11796676 |
| Partition coefficient (Pliska et al., 1981) | 88,72 | 15609 | 91,31 | 5959 | 80,70 | 64864 | -21,55 | 22859192 | 17,93 | 3083900 |
| The stability scale from the knowledge-based atom-atom potential (Zhou-Zhou, 2004) | 88,12 | 14757 | 86,38 | 7958 | 80,00 | 47212 | -13,90 | 16850368 | 25,19 | 4463524 |
| TOTLS index (Cornette et al., 1987) | 87,55 | 45031 | 90,38 | 22059 | 86,10 | 44840 | -21,99 | 22607776 | 8,52 | 6911906 |
| Modified Kyte-Doolittle hydrophobicity scale (Juretic et al., 1998) | 87,49 | 30532 | 89,82 | 18578 | 91,35 | 27872 | -27,69 | 22955832 | 8,04 | 7978956 |
| PRIFT index (Cornette et al., 1987) | 87,33 | 24215 | 93,47 | 7791 | 91,17 | 5424 | -24,24 | 23311728 | -0,16 | 10847386 |
| Average gain ratio in surrounding hydrophobicity (Ponnuswamy et al., 1980) | 87,27 | 38770 | 85,30 | 44262 | 76,86 | 119120 | -19,54 | 20484840 | 17,42 | 3914864 |
| AA composition of MEM of multi-spanning proteins (Nakashima-Nishikawa, 1992) | 87,08 | 60266 | 89,87 | 56417 | 91,92 | 91088 | -27,23 | 20384792 | 1,39 | 11341172 |

**Table 4.** The 20 properties with the highest optimization percentages from the AAindex database (Kawashima and Kanehisa, 2000). Optimality of the standard genetic code in the coevolution theory framework. For each global index of spatial autocorrelation, two different optimality measures of the genetic code are reported in the restricted set of permutation codes subject to biosynthetic constraints. The first, Opt, represents the optimization percentage (see Materials and methods) and the second, Nsg, indicates the number of codes that have the relative index value strictly greater than that of the genetic code. The listed properties have the highest optimization percentages, in relation to index I, among all those of the database. All spatial weight matrices here are row-standardized. The italic writing emphasizes that the property was added for comparison to the AAindex database.

| Global spatial autocorrelation indices<br>(related to different spatial row-standardized weighting schemes) | I |  | I<sub>C</sub> |  | I<sub>1</sub> |  | I<sub>2</sub> |  | I<sub>R</sub> |  |
| --- | --- | --- | --- | --- | --- | --- | --- | --- | --- | --- |
| Optimality measures | Opt | Nsg | Opt | Nsg | Opt | Nsg | Opt | Nsg | Opt | Nsg |
| Conformational preference for parallel beta-strands (Lifson-Sander, 1979) | 96,16 | 12466 | 88,92 | 84706 | 86,67 | 183044 | -22,39 | 19203776 | 18,46 | 5551144 |
| Relative partition energies derived by the Bethe approximation (Miyazawa-Jernigan, 1999) | 95,93 | 2740 | 97,77 | 505 | 93,18 | 6156 | -23,03 | 21162128 | 31,50 | 1844060 |
| *Effective mean energy (Miyazawa-Jernigan, 1996)* | 95,79 | 2727 | 97,71 | 523 | 92,97 | 6448 | -23,25 | 21183384 | 32,01 | 1804060 |
| Optimized relative partition energies - method A (Miyazawa-Jernigan, 1999) | 95,78 | 2800 | 97,66 | 549 | 92,98 | 6288 | -23,07 | 21173416 | 31,07 | 1886024 |
| Buriability (Zhou-Zhou, 2004) | 92,64 | 3232 | 94,11 | 2488 | 73,09 | 89144 | -11,80 | 16787312 | 45,72 | 868924 |
| Interactivity scale obtained by maximizing the mean of correlation coefficient over single-domain globular proteins (Bastolla et al., 2005) | 92,53 | 4820 | 92,78 | 4106 | 74,73 | 70832 | -9,42 | 16214376 | 31,19 | 1824056 |
| Effective partition energy (Miyazawa-Jernigan, 1985) | 92,48 | 9132 | 95,71 | 2831 | 89,50 | 22952 | -28,33 | 21768312 | 43,01 | 932600 |
| HPLC parameter (Parker et al., 1986) | 91,89 | 5058 | 94,83 | 798 | 84,01 | 18512 | -16,94 | 19212280 | 26,77 | 2165220 |
| PRILS index (Cornette et al., 1987) | 91,39 | 7900 | 92,22 | 11530 | 89,72 | 9364 | -22,26 | 21169208 | 14,94 | 4783316 |
| Polarity (Grantham, 1974) | 90,21 | 4180 | 92,31 | 2366 | 77,72 | 60768 | -26,79 | 22064000 | 36,37 | 1341884 |
| Hydropathy index (Kyte-Doolittle, 1982) | 89,92 | 22172 | 91,76 | 14196 | 89,28 | 17396 | -32,05 | 24202720 | 14,34 | 5565168 |
| Surface composition of amino acids in extracellular proteins of mesophiles (percent) (Fukuchi-Nishikawa, 2001) | 89,82 | 4330 | 92,16 | 11515 | 51,94 | 474528 | 1,43 | 11799184 | 49,46 | 464724 |
| Optimized relative partition energies - method B (Miyazawa-Jernigan, 1999) | 89,43 | 4451 | 90,49 | 3520 | 80,20 | 25872 | -20,11 | 20469000 | 27,67 | 2051100 |
| Optimized relative partition energies - method C (Miyazawa-Jernigan, 1999) | 89,35 | 3433 | 92,22 | 1508 | 79,69 | 22840 | -20,32 | 20088272 | 32,39 | 1693868 |
| Modified Kyte-Doolittle hydrophobicity scale (Juretic et al., 1998) | 88,76 | 27850 | 88,84 | 24128 | 86,83 | 32888 | -28,13 | 23122312 | 15,26 | 5435068 |
| van der Waals parameter R0 (Levitt, 1976) | 88,48 | 28800 | 68,58 | 142848 | 62,51 | 408672 | 16,05 | 6689088 | 44,75 | 1356480 |
| Average surrounding hydrophobicity (Manavalan-Ponnuswamy, 1978) | 88,47 | 56666 | 91,11 | 17802 | 86,55 | 84188 | -32,24 | 24044296 | 9,14 | 7332424 |
| Interactivity scale obtained from the contact matrix (Bastolla et al., 2005) | 88,40 | 18155 | 94,44 | 6311 | 85,05 | 60686 | -20,87 | 21132904 | 9,42 | 6744660 |
| *Updated Polar Requirements (Mathew and Luthey-Schulten, 2008)* | 88,38 | 1054 | 91,82 | 2484 | 69,64 | 157632 | -5,68 | 14479536 | 70,39 | 68192 |
| AA composition of MEM of multi-spanning proteins (Nakashima-Nishikawa, 1992) | 88,25 | 37920 | 92,24 | 26450 | 91,23 | 37684 | -30,48 | 22652968 | 5,03 | 9304112 |

Since the first step (on which the choice of the first 50 properties with the highest estimates depends) is based on sampling, there could be doubts (even with a large sample of codes, equal to 1/50 of the population size) that could be expected some uncertainty in the results, in relation to which properties will appear in the final tables. To dispel any doubts, we exhaustively computed the optimization percentage of the top 100 properties with the highest estimates, confirming the stability of the tables. In conclusion, Tabs. 3 and 4 actually report the 20 properties with the highest optimization percentages, relating to index I, among all the properties analyzed in the database.

It turns out that the only properties able of exceeding the 95% threshold in the optimization percentage relative to index I in both Tabs. 3 and 4 are the partition energies of the amino acids and the conformational preference for parallel beta-strands. Therefore, it is difficult to escape the conclusion that they played the key role in the origin of the organization of the genetic code.

Furthermore, three measures of the partition energy of amino acids - highly correlated with each other - show on the columns of the genetic code (index I_C_) an optimization percentage of about 98% occupying the first three ranks in both Tabs. 3 (without row-standardizing) and 4 (with row-standardized spatial weight matrices).

Conversely, operating with non-row-standardized weight matrices, no measure of amino acid polarity is present within the first twenty more optimized properties (Tab. 3). However, we had the opportunity to verify that the first polarity measure to appear is that of Grantham (1974) with only 85.51% of optimization; if, however, we restrict the analysis to only the columns of the genetic code (Ic index) it manages to achieve a good level of optimization of 95.68%. However, it is appropriate to specify that it is the genetic code in its entirety that is subjected to the scrutiny of natural selection, therefore it is the first percentage that is truly relevant, that is, the one referred to the changes involving the whole code and not just a part of it, as happens for column optimization. It follows that, with optimization percentages of 85.51% or lower, it is reasonable to believe that polarity might not have played a key role in the origin of the organization of the genetic code. Optimization levels lower than 90% are, in actual fact, completely common among the physicochemical properties of amino acids (Tabs. 3 and 4) so this behavior could simply reflect the strong correlation between the polarity measurements and those of the partition energies of the amino acids.

Standardization, on the other hand, seems to somehow favor polarity, but the level of optimization achieved is still quite low (90.21% for the I index and 92.31% for Ic); however, various polarity measures are present in the tables constructed using standardized weights (four measures in Tab. 2 and two in Tab. 4). However, we believe that the natural condition is better preserved by operating using the weights provided directly by the structure of the genetic code, i.e. without row-standardizing the spatial weight matrices (Tabs. 1 and 3), as opposed to standardization which inevitably destroys their symmetry.

Another observation in favor of polarity is that concerning the good performance of the Updated Polar Requirements (Mathew and Luthey-Schulten, 2008) in achieving the lowest value among those recorded in the second column of Tab. 4. It refers to the number of codes with strictly better values of the Moran’s I than the one achieved by the genetic code. It should be noted, however, that the relevance of the result is reduced by the fact that the corresponding optimization percentage is only about 88% (Tab. 4).

### The codes that perform better than the genetic code and their relationships with the latter: the genetic code is neither a global nor a local maximum, also there is a directed path by the genetic code to a code with a value of Moran I close to the maximum absolute value

Referring to the global Moran’s I index of spatial autocorrelation (Tab. 3) and relative partition energies derived by the Bethe approximation (Miyazawa-Jernigan, 1999), Fig. 1 plots the histogram and Fig. 2 reports the permutation of amino acids within the code table that maximizes the value of the index (I_max_). As you can see the genetic code differs from this code with maximum value of Moran I for the different encoding of seven blocks of synonymous codons. In particular, in order to move from the genetic code to the code of Fig. 2b, a number of base substitutions should be made. In a sense, this indicates a considerable mutational distance because the (Asp Thr Lys) 3-cycle and the (Glu Pro) transposition (Kirson, 2017) imply multiple mutational events, while still occurring between amino acids belonging to the same biosynthetic class. It should also be noted that all these seven amino acid changes (Fig. 2) involve only amino acids of the biosynthetic families of Asp and Glu and that - at least at certain points in the evolutionary history of the genetic code - they might have taken place, precisely because of the existence of the mechanism that allowed amino acids belonging to the same biosynthetic family to be distributed on contiguous codons (Di Giulio, 2002, 2019). However, this probably did not happen due to the presence of strong historical constraints on the Asp and Glu codons, which were among the first amino acids to be encoded in the genetic code (Ikehara et al., 2002; Di Giulio, 2008a). The (Arg Gln) transposition might also have been hampered because Arg appears to have had strong physicochemical constraints due to its ability to stabilize the structure of proteins at high temperature and high pressure (Di Giulio, 2000a, 2005a, 2005b, 2005c). Such physicochemical constraints would have required a high number of codons to code for this amino acid in the genetic code (Di Giulio, 2000a, 2005a, 2005b, 2005c). If this was the case, then the organization of the structure of the genetic code would be close to that of a global optimum more than the optimization percentages reported in Tabs. 3 and 4 would indicate.

**Figure 1.**
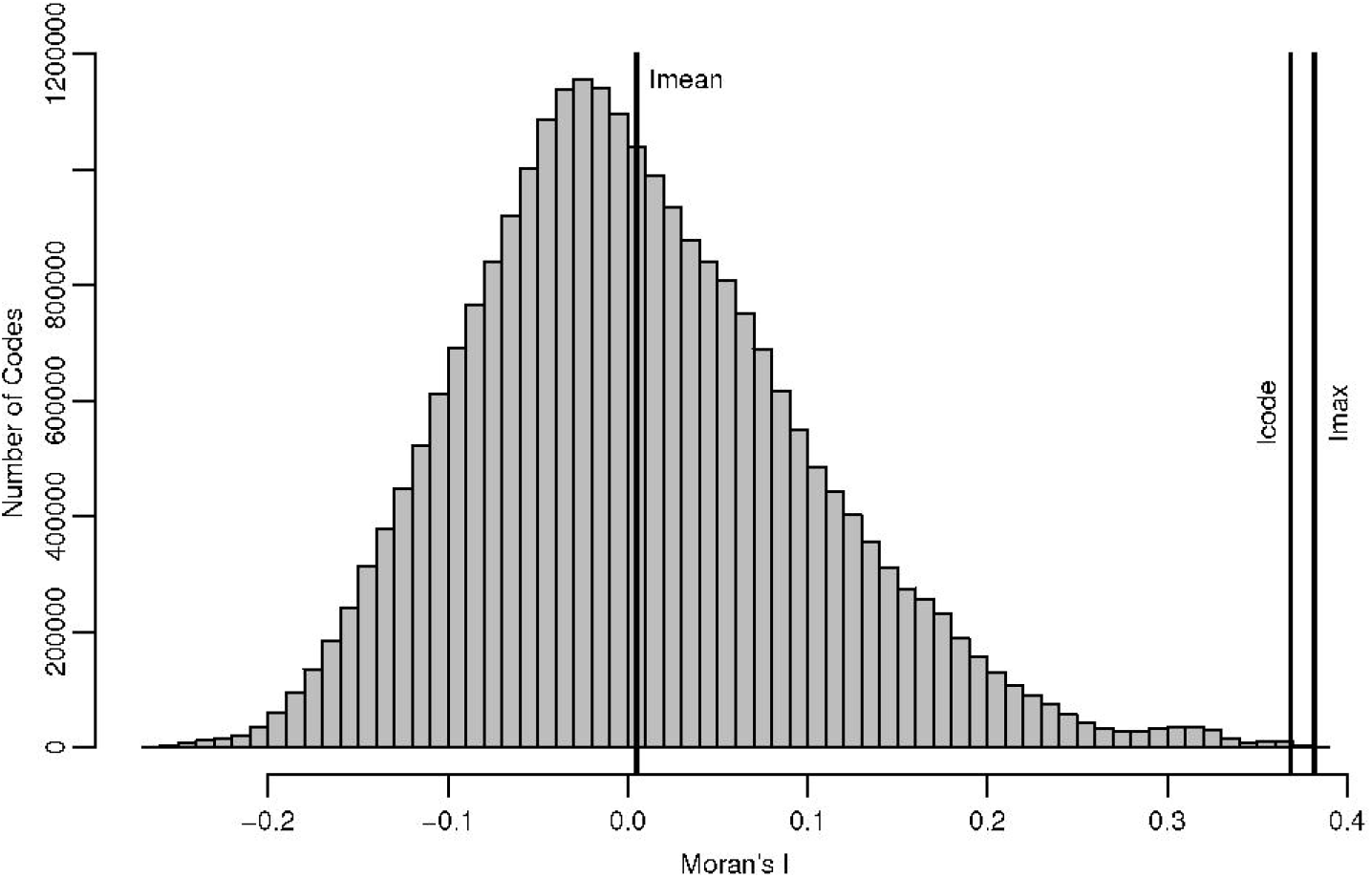
Histogram for the Moran’s I values obtained from all the 24,883,200 possible permutations of the genetic code generated by amino acid restricted permutations subject to the biosynthetic constraints and referring to relative partition energies derived by the Bethe approximation (Miyazawa-Jernigan, 1999). The three vertical lines indicate the mean value (I_mean_ = 0.00497), the maximum (I_max_ = 0.38190) and the value of Moran’s I associated with the genetic code (I_code_ = 0.36853). Therefore, the optimization percentage (see text) reached by the genetic code is 96.45% (see Tab. 3). Note that here the spatial weight matrix for the calculation of Moran’s I is not row-standardized.

**Figure 2.**
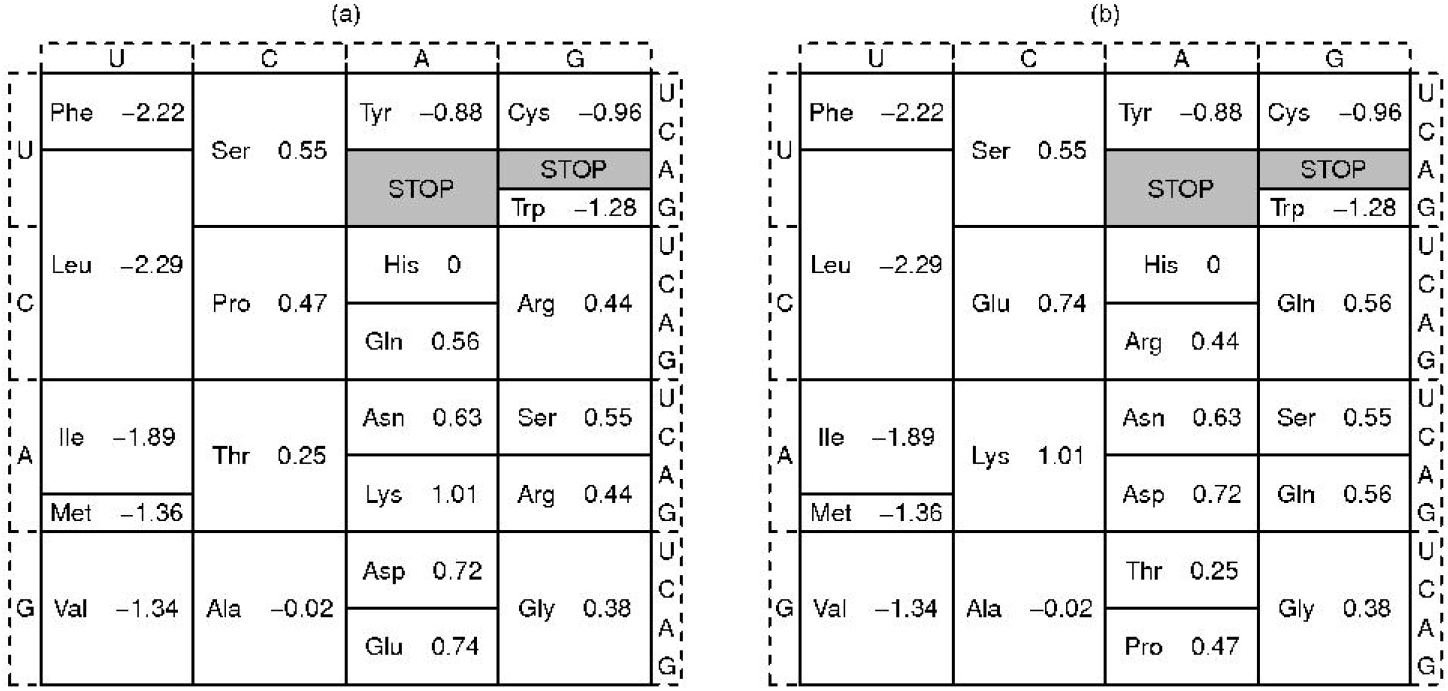
Comparison between the natural genetic code (a) and the table of the code (b) that maximizes the Moran’s I index of spatial autocorrelation in the restricted permutations set. The permutation of amino acids among synonymous codon blocks that changes (a) into (b) can be expressed as the product of three disjoint cycles: (Arg Gln) ◦ (Glu Pro) ◦ (Asp Thr Lys). In each block of synonymous codons, in addition to the coded amino acid, the corresponding value of the relative partition energies derived from the Bethe approximation (Miyazawa-Jernigan, 1999) is also reported.

The genetic code is not a local maximum. Indeed, of the 190 possible unordered amino acid pairs, only 35 are those with both amino acids belonging to the same biosynthetic family. Therefore, in the restricted set of amino acid permutations subject to biosynthetic constraints, only 35 are those that differ by a single amino acid exchange from that of the genetic code. Choosing as distance between permutations (and between corresponding codes) an edit-distance counting how many interchanges (i.e., transpositions of two arbitrary amino acids) have to be performed to transform one permutation into another, these 35 permutations are the only ones to have not null minimal distance from the genetic code. Now, for the genetic code to be a local maximum, all of these 35 permutations must have a Moran’s I value no higher than the value of the code itself (I_code_). Instead, some of these [precisely: (Pro Arg), (Gly Ser), (His Pro), (Asp Asn), (Asn Thr), (Asp Thr) and (Glu Pro)] have a Moran’s I value greater than I_code_, so it follows that the genetic code is not a local maximum. For example, an exchange between Asn and Thr (which differ by a single base in some of their codons) is enough to achieve an improvement in the Moran’s I corresponding to an increase of 1.16% in the optimization percentage. All other transpositions listed have lower increments, except (Asp Thr) (an increase of 2.46%), in this case, however, their codons differ by at least two bases.

Continuing to refer to the relative partition energies derived by the Bethe approximation (Miyazawa-Jernigan, 1999), we find that in the restricted set of 24,883,200 codes obtained from amino acid permutations subject to biosynthetic constraints there are 4429 codes with a Moran I strictly greater than the natural code (Table 3). Defining adjacent two of these (4429+1) codes if and only if they differ by a single transposition of two amino acids encoded by (at least some) codons differing by only a single nucleotide; it is of some interest to verify the reachability of the code with the maximum value of Moran I (Fig. 2) - starting from the natural code - through a directed path in which each step is associated with a non-negative [or even slightly negative (≥ −0.0005)] variation of Moran’s I. Well, under these conditions it is shown that such a path does not exist. However, another code with an index value (I’=0.38009) very close to the global maximum (I_max_=0.38190) is accessible, which can be obtained from the natural code through the following sequence of transpositions: (Arg Pro), (Asp Asn), (Lys Thr), (Gln Arg), (Asp Thr), and (Asn Thr). Therefore, in recalculating the optimization percentage (being invariant I_code_=0.36853 and I_mean_=0.00497) instead of I_max_=0.38190 we should consider I’_max_=0.38009, thus recording a slight increase from 96.45% to 96.92% of the optimization percentage. In other words, even if under quite biologically reasonable conditions, the code with the highest value of Moran I is not actually achievable, nevertheless the optimization percentage is not affected when it is referred to the maximum value that is actually achievable. In short based on these assumptions, the reachability or not of the code with the maximum value of Moran’s I does not significantly affect the conclusions.

## Discussion

### The partition energy of amino acids: its importance in the origin of the genetic code and implications for the theories proposed to explain this origin

Some considerations would seem to be highly sustainable based on the results reported here. The first is that the partition energy (Miyazawa and Jernigan, 1985, 1996, 1999) is the amino acid property that seems to be better reflected in the organization of the genetic code and not the polarity of amino acids (Tabs. 1, 2, 3 and 4), as instead widely supported and assumed in literature (Woese, 1965, 1967; Woese et al., 1966; Di Giulio, 1989a; Haig and Hurst, 1991; Freeland and Hurst, 1998; Butler et al., 2009; Buhrman et al., 2013; Błażej et al., 2016, 2018, 2019; Wnętrzak et al., 2018). This consideration would be consistent with a vision of the origin of the genetic code that focuses on enzymatic catalysis as the main selective pressure that would have triggered its origin (Wong, 1976, 1991, 1980; Di Giulio, 1997a, 2003, 2008b, 2015). Indeed, the partition energy of amino acids would be closely related to the structure of proteins (Guy, 1985; Miyazawa and Jernigan, 1985, 1996) which in turn would reflect the enzymatic catalysis. In extreme synthesis, Miyazawa and Jernigan (1985, 1996, 1999) inter-residue contact energies were extracted, using methods of statistical mechanics, from observed contact frequencies of residue pairs in known crystalline structures of globular proteins and are generally considered a good approximation of realistic protein interactions. Based on these data, the partition energies were also derived for the 20 amino acids, which are related to the propensity of residues to be buried in the interior of proteins or to be exposed to water on the surface (Miyazawa and Jernigan, 1985, 1996). Now, the finding that precisely the amino acid properties so exquisitely related to the protein structure as the partition energies are also the best autocorrelated in the structure of the genetic code (Tabs. 1 and 2) and the best optimized there (Tabs. 3 and 4), suggests that the structuring itself of the genetic code might have been to a large extent oriented by the need to preserve protein structure and its basic functions, first of all enzymatic catalysis, and to promote its evolutionary improvement.

On the contrary, the low performances of other properties, in particular of polarity, which refer to stereochemical interactions between amino acids and codons or anticodons, indicate that physicochemical properties found in the organization of the genetic code (Woese et al., 1966; Alff-Steinberger, 1969; Jungck, 1978; Nelsestuen, 1978; Wolfenden et al., 1979; Sjostrom and World, 1985; Di Giulio, 1989a, 1989b, 1991, 2017a, 2018b; Chiusano et al., 2000; Lacey Jr. et al., 1992; Freeland et al., 2000; Buhrman et al., 2013; Błażej et al., 2016, 2018, 2019; Wnętrzak et al., 2018) might more likely be considered expressions of the enzyme catalysis itself rather than of presumed interactions between amino acids and codons or anticodons. The whole argument appears more easily interpretable within the coevolution theory of the genetic code (Wong, 1976, 1980; Di Giulio, 2018a) than within the physicochemical theories (Sonneborn, 1965; Woese, 1965, 1967; Lacey Jr et al., 1992; Fitch and Upper, 1987; Higgs, 2009). Indeed, for the coevolution theory, enzymatic catalysis would have been the main selective pressure that would have triggered the origin of the genetic code (Wong, 1976, 1980; Di Giulio, 1997a, 2003, 2008b, 2015, 2018a). On the contrary, physicochemical theories, in general, argue that the optimization of physicochemical properties detectable in the organization of the genetic code (Tabs. 3, and 4) is an expression of interactions between amino acids and codons or anticodons that would have promoted his origin (Woese, 1967; Mathew and Luthey-Schulten, 2008). In other words, our results (Tabs. 1, 2, 3 and 4) are easily explained by the coevolution theory of the genetic code (Wong, 1975; Di Giulio, 2008a, 2016a, 2016b, 2017a, 2017b) because it argues that during the origin of the genetic code the biosynthetic relationships between amino acids conditioned above all the evolution of its rows (Taylor and Coates, 1989; Di Giulio, 2001b, 2008a, 2017b, 2018b, 2019), and therefore on the columns of the code might be allocated as best as possible amino acids with similar physicochemical properties (Wong, 1980; Di Giulio, 2017b, 2018a, 2019), as indeed it has been observed (Nelsestuen, 1978; Wolfenden et al., 1979; Sjostrom and World, 1985; Di Giulio, 1989b, 2017a, 2018b; Chiusano et al., 2000). On the contrary, the physicochemical theories (Sonneborn, 1965; Woese, 1965, 1967; Lacey Jr. et al., 1992; Fitch and Upper, 1987; Higgs, 2009) would not be able to give an equally satisfactory description of these observations (Di Giulio, 2016a, 2016b, 2017a, 2017b), also because they would be unable to explain why the physicochemical properties of amino acids do not show a significant distribution on the rows of the code (Di Giulio, 2016a, 2016b, 2017a, 2017b), although some significance is observed for the polarity of amino acids (P = 0.016 (1548/10^5^); Tab. 2). Furthermore, while the coevolution theory would seem to be able to explain the statistical significance of the distribution linked both to the biosynthetic relationships of amino acids and to their physicochemical properties, the physicochemical theories would not have this capacity (Di Giulio, 2017b).

The very high optimization of the partition energy of amino acids, in particular on the columns of the genetic code (Tabs. 3 and 4), is in agreement with the hypothesis of Woese (1965, 1967) that one of the main selective pressures that would have structured the genetic code would have been the lowering of translational noise. Other forms of reduction of ambiguity hypothesized to be present in some stages of the origin of the genetic code (Fitch and Upper, 1987; Ardell and Sella, 2001; Barbieri, 2015), would also be compatible with the very high optimization found (Tabs. 3 and 4). On the other hand, any theory that is not capable of predicting a suitable level of optimization would be falsified by the evidence reported here. Stereochemical theory (Woese, 1967; Shimizu, 1982; Szathmáry, 1993; Yarus, 2017) might be one of the theories unable to make this prediction. Indeed, the very high optimization would seem more the consequence of the lowering of the translation noise through natural selection than of interactions, for example, between amino acids and anticodons. Precisely, these interactions might not have been able to determine such different behavior between the columns and rows of the genetic code as regards their level of optimization (Tabs. 3 and 4). In other words, if the very high optimization (Tabs. 3 and 4) had really been the result of interactions between amino acids and codons or anticodons then we should have observed a less marked behavior in favor of column optimization and against that of row because it is the first two positions of the codons that determine most of the meaning of a certain codon. Whereas, in the event that it was only the presence of the second base of the codon or anticodon that was important in this stereochemical interaction - which hypothetically could have promoted the origin of the genetic code (Weber and Lacey, 1978; Lacey Jr. et al., 1992) - then this interaction involving a single base might not have ensured the correspondence between a determined anticodon (or a codon) and a specific amino acid, as is on the contrary maintained by the stereochemical theory as its main assumption. Therefore, in this sense, the stereochemical theory would not be corroborated by the very high optimization of the column found (Tabs. 3 and 4), rather it would be falsified.

In general terms, the observations reported here are at least compatible with the theory of the four columns of Higgs (2009) because the very high optimization that has been achieved on the columns (Tabs. 3 and 4) would evidently corroborate this theory, as its name suggests. Nevertheless, entering specifically, the model of evolution of the genetic code analyzed here incorporates the biosynthetic constraints derived from the coevolution theory of the genetic code (Facchiano and Di Giulio, 2018) which are absolutely neither foreseen nor compatible with the theory of the four columns. Indeed, this suggests that the organization of the genetic code occurred by allocating amino acids with similar physicochemical properties along the columns of the genetic code even if the driving force during this origin was not the minimization of translational error, but positive selection for the increased diversity and functionality of proteins (Higgs, 2009). On the contrary, the coevolution theory would be able to explain the very high optimization observed on the columns (Tabs. 3 and 4) simply by assuming that while the code was originating along its rows (Di Giulio, 2008a, 2017b, 2019), its columns were allocated similar amino acids to reduce, for example, the effects of translation errors (Wong, 1980; Di Giulio, 2017b, 2018a), i.e. by incorporating an important selective pressure envisaged by the physicochemical postulates (Woese, 1965, 1967). That is to say, the coevolution theory would be highly compatible with the very high observed column optimization (Tabs. 3 and 4; Facchiano and Di Giulio, 2018). This would not be true for the theory of the four columns which (i) would not be able to explain an aspect incorporated in the model used here, namely the high statistical significance of the distribution of the biosynthetic pathways on the lines of the genetic code (Taylor and Coates, 1989; Di Giulio, 2008a, 2017b, 2018b; Di Giulio and Amato, 2009) simply because this theory would suggest nothing regarding the biosynthetic pathways of amino acids (Higgs, 2009), even if (ii) it would explain the very high column optimization (Tabs. 3 and 4) but not through a lowering of the translation noise and ambiguity present in the primitive protocell but through a positive selection to increase the diversity and functionality of proteins, as claimed by Higgs (2009). To us, however, it seems no less reasonable that if the aim had been to promote the diversity and functionality of the proteins of the evolving genetic code (Higgs, 2009) - i.e. ultimately the structure of proteins - then this would have been more easily achieved by lowering the translational noise and ambiguity present in the primitive protocell. Furthermore, it is likely that this very high optimization reflects a high rate of adaptive evolution which might in actual fact have been highest among amino acids that are more similar (Bergman and Eyre-Walker, 2019). All this would, however, have allowed a better maintenance of the protein structure through the generations, that is to say, a better preservation of the enzymatic catalysis over time, and would have favored its evolution. In conclusion, the four-column theory could have substantially correct aspects but would only take care of the aspect related to the selective pressure that would have triggered the origin of the genetic code, not being able, in our opinion, to actually explain the fundamental mechanism through which amino acids were allocated in the genetic code which surely depended on their biosynthetic pathways given the presence of the molecular fossils of these pathways (Wong, 1976, 1988; Wachtershauser, 1988; Danchin, 1989; de Duve, 1991; Edwards, 1996; Di Giulio, 1992, 1997b, 2002, 2008a).

### The β-sheets and β-strands of proteins, the biosynthetic relationships between amino acids and the origin of the organization of the genetic code

One of the main predictions of the theory of coevolution is that some of the fundamental themes of the structure of proteins are present in the organization of the genetic code because the main selective pressure tending to organize the code should have been, in the last analysis, the very structure of proteins (Di Giulio, 1996). In this regard, Orgel (1977) discussed the importance of β-turns stabilized by β-sheets as plausible sites of the early enzymatic activity (see also, Kun et al., 2008). Brack and Orgel (1975) suggested that the earliest form of genetic coding might have specified polypeptides with a strong tendency to form stable β−sheet structures. Moreover, Jurka and Smith (1987) suggested that β−turns became objects of selection in the prebiotic environment and affected the evolution of the genetic code and the biosynthetic pathways of amino acids (see also, Kun et al., 2008). It has also been suggested that β−sheets are related to amino acid pairs in a precursor-product biosynthetic relationship (Di Giulio, 1996), i.e. that the biosynthetic pathways of amino acids were also selected because they reflect β−sheets (Di Giulio, 1996).

An observation compatible with these suggestions was reported here. In fact, it has been observed that the conformational preference for parallel β-strands (Lifson and Sander, 1979) show very high optimization percentages (of 95.45% and 96.16% respectively in Tabs. 3 and 4). It is therefore possible to hypothesize that even the β−strands might have become the object of selection during the origin of the genetic code and might have conditioned the choice of biosynthetic relationships between amino acids. In other words, the biosynthetic relationships between the amino acids were also chosen on the basis of the similarity of their conformational preferences for β−strands, as corroborated by their high optimization percentages (Tabs. 3 and 4). This would be compatible with the observation that β−sheets seem to have conditioned the choice of amino acid pairs that are in biosynthetic precursor-product relationship (Di Giulio, 1996). Ultimately, the theory of coevolution would be corroborated because some of the fundamental themes of the structure of proteins, such as β−sheets and β−strands, would be reflected in the organization of the genetic code.

### The coevolution theory of the genetic code origin, its evolutionary contingency and the almost global optimality of the code

The coevolution theory of the genetic code is undoubtedly a theory of a nondeterministic nature in the sense that if the evolution of the genetic code started all over again, then we could in all probability observe a genetic code completely different from the current one, even assuming that its origin was once again influenced by the biosynthetic relationships between amino acids. This is because the biosynthetic relationships between amino acids would not be constrained at all or at least marginally and this would obviously depend on the fact that a given biosynthetic pathway would evolve along evolutionary paths that are also completely different from the chemical point of view of its intermediate and final products. Therefore, the coevolution theory would imply (or at least it would not exclude) that a new evolution ab initio of the genetic code might lead to the encoding of amino acids even completely different from those observed today (Wong, 1975; Di Giulio, 2008a). On the contrary, stereochemical theory would not admit such a prediction because, being a theory of a deterministic nature, it would impose that a new evolution of the genetic code could only re-determine the same type of genetic code (Crick, 1968), at least in one of its more radical interpretations. This being the case, it could be at least curious that the genetic code has reached almost an overall optimum, given that this very high optimization would seem more typical of a physicochemical process than of a biological process such as that of the origin of the genetic code envisaged by the coevolution theory (Wong, 1975; Di Giulio, 2008a). In other words, it seems surprising that - under the model predicted by the coevolution theory - it might nevertheless have been observed that the genetic code is close to an overall optimum (Tabs. 3 and 4); this theory, in actual fact, would seem to be linked to a high degree of evolutionary contingency which at first sight appears incompatible with this level of optimization. A possible explanation is that this level of optimization would have been achieved by natural selection by exploiting precisely the fundamental mechanism of structuring the genetic code contemplated by the coevolution theory, in order to almost optimally allocate amino acids with similar physicochemical properties on the columns of the genetic code (Wong, 1980; Di Giulio, 2018a). That is to say, natural selection, working on the structuring of a genetic code that was evolving through a general plan of evolutionary contingency (Blount, 2016) imposed by the biosynthetic relationships between amino acids (Wong, 1975; Di Giulio, 2008a), would have managed, despite this, to bring the code towards this almost global optimality (Tabs. 3 and 4) under the very strong selective pressure aimed at perpetuating the enzymatic catalysis through the preservation of the protein structure. In other words, two elements would have played a key role in the origin of the organization of the genetic code: one linked to the contingency of the biosynthetic relationships between amino acids and the other represented by the need to optimize those properties (physicochemical or biological) truly decisive for the perpetuation of enzymatic catalysis and for its evolutionary improvement. In summary, the evolutionary contingency linked to the way of operating typical of the coevolution theory might have led to a genetic code coding for amino acids even completely different from those we are used to (Stegmann, 2004), but, nevertheless, the level of optimization of the table of this new code could inevitably have been very close to what we have observed here.

### An important implication: error minimization is a property of the genetic code that emerged as a result of natural selection and is not instead a neutral property, that is, originated as a by-product of another mechanism

The observations presented here (Tabs. 1, 2, 3 and 4) have a direct relevance to understand whether the so-called error minimization possessed by the genetic code is a property that emerged as a result of natural selection (Di Giulio, 2000b, 2018a; Stoltzfus and Yampolsky, 2007) or if on the contrary it has not been selected but is an emergent property, i.e. neutrally originated, as a consequence of other mechanisms that were structuring the genetic code (Massey, 2008, 2016, 2018, 2019; Koonin, 2017). Given that a model has been investigated here that combines two factors believed to be responsible for the origin of the genetic code - that is to say, the biosynthetic relationships between amino acids and their physicochemical properties (Di Giulio, 2013) - we are in a position to respond to this question at least under these conditions. Indeed, the mechanism on which the coevolution theory is based (Wong, 1975; Wachtershauser, 1988; Danchin, 1989; de Duve, 1991; Edwards, 1996; Di Giulio, 1997b, 2002, 2004, 2008a) could actually have been the mechanism that was structuring the genetic code (Di Giulio, 2018a) and therefore - in accordance with the neutral hypothesis (Massey, 2008, 2016, 2018, 2019) - might have given rise to the minimization of the error as its by-product, without direct action of natural selection (Massey, 2008, 2016). It would seem self-evident that the levels of optimization reached by the genetic code with the relative probabilities (Tabs. 3 and 4) might not be compatible with a neutral origin because, for example, it would be unlikely that a neutral mechanism was able to generate optimization percentages close to 100% (Tabs. 3 and 4). In other words, it appears statistically very unlikely that a neutral mechanism might have been responsible for this very high optimization (Tabs. 3 and 4) because if the near-optimality of the genetic code had been a by-product of another mechanism, then this would have probably had to produce much lower optimization percentages and far from those reported in Tabs. 3 and 4, precisely because they were not directly selected. In particular, it is easy to prove, whatever the amino acid property considered, that the average value of the optimization percentages of the 24,883,200 codes subject to biosynthetic constraints is zero (results not shown). Then, under the neutral hypothesis, we should have observed an optimization percentage of the genetic code not significantly far from that expected, that is, from zero. In other words, the biosynthetic pathways structuring, by hypothesis, the genetic code might not have determined - as an emergent property - an optimization percentage significantly different from the null average value. Given that, on the other hand, the percentage of optimization found, for example, for the relative partition energies derived by the Bethe approximation is of 95.93% (P=1.10×10^-4^) (Tab. 4), the hypothesis that the near-optimality of the genetic code arose entirely neutrally, i.e. without any action of natural selection, can be rejected. Thus, at least with reference to the model analyzed here, the neutralist hypothesis is unable to explain the observed high level of the optimization percentage, which is instead expected under the selectionist hypothesis (Di Giulio, 2018a).

Finally, with reference to this same property, the comparison between row and column optimizations is extremely instructive. Indeed, if we restrict the analysis to just the rows (i.e. if we consider the I_R_ index) - on which the biosynthetic relationships between amino acids are mostly distributed (Di Giulio, 2008a, 2016b, 2017a, 2018b) - we find an optimization of 31.50% (P = 0.074) (Tab. 4). Such a value of the optimization percentage might be considered congruous to that expected, under the neutralist hypothesis, if we consider that the biosynthetic pathways connecting the amino acids are structuring the genetic code as maintained by the coevolution theory. However, given that on the columns of the genetic code we observed an optimization percentage of 97.77% (P= 2.0×10^-5^) (Tab. 4), we must conclude that there was the intervention of another force, namely natural selection, precisely because the distribution of biosynthetic relationships between amino acids is not very significant on the columns (Di Giulio, 2008a, 2016b, 2017a, 2018b). More generally, the fact that optimization is mainly linked to the columns of the genetic code (Tabs. 1, 2, 3, and 4) would seem incompatible with a neutral process, because if this had really been operational then it would not have been able to make a so marked distinction between the columns and rows of the genetic code as, for example, the one reported in Tabs. 3 and 4 (see also Di Giulio, 2016b, 2017a, 2018b).

In conclusion, the neutral hypothesis of the origin of error minimization (Massey, 2008, 2016, 2018, 2019; Koonin, 2017) should have been falsified under the hypothesis that the genetic code was structured using the mechanism envisaged by the coevolution theory.

Koonin (2017) also argues that error minimization, despite being a key property of the genetic code, “likely evolved as a by-product of code expansion rather than by direct selection for code robustness” because “the coevolution model, although producing some level of error minimization, was substantially inferior to the other scenarios “(Koonin, 2017). Koonin (2017) seems to intend that the coevolution theory is not satisfactory because in the simulation models of Massey (2008, 2016) it fails to provide levels of error minimization comparable to those of other scenarios used in these simulations. Now, the fact that the coevolution theory fails to provide certain levels of error minimization (Massey, 2008, 2016; Di Giulio, 2018a) would not mean that this theory might be inconsistent - as Koonin (2017) would seem to understand - but that the neutral hypothesis of error minimization can instead be (Di Giulio, 2018a). Indeed, the results reported here would seem to prove just that, because such high optimization percentages and probabilities so far from random (Tabs. 3 and 4) - obtained under the model of coevolution theory - would be able to show how the genetic code is positioned - in the set of amino acid permutation codes subject to biosynthetic constraints (Facchiano and Di Giulio, 2018) - in extreme optimization regions (Fig. 1) such that they cannot be reached through a simple emergent and therefore neutral process, but only through a mechanism guided by natural selection.

In conclusion, we must believe that the extraordinary property of error minimization possessed by the genetic code did not emerge as a consequence of a simple neutral process - as claimed by Massey (2008, 2016, 2018, 2019) and Koonin (2017) - but is instead a clear manifestation of the strength of natural selection (Di Giulio, 2018a).

## Conclusions

We believe that the results reported here (Tabs. 1, 2, 3 and 4) make the following conclusions very plausible: (i) that partition energy is really the amino acid property that can provide us with the key to understanding the origin of organization of the genetic code; (ii) that the polarity of the amino acids was not instead taken into account during this origin; (iii) that the allocations of amino acids within the blocks of synonymous codons of the respective biosynthetic families were such as to make the genetic code close to a global optimum; (iv) that the near-optimality of the genetic code has emerged, at least for the model analyzed here, by the action of natural selection; and (v) that the conformational preference for parallel β-strands, conditioning the choice of biosynthetic relationships between amino acids, played a not secondary role in the organization of the genetic code. Finally, we are convinced that all the observations reported here are highly compatible with the coevolution theory of the origin of the genetic code (Wong, 1975; Di Giulio, 2008a, 2016a, 2016b, 2017a, 2017b) which we therefore believe to be a highly corroborated theory (Wong, 2007; Di Giulio, 2009), and that, on the contrary, the physicochemical theories are not (Sonneborn, 1965; Woese, 1965, 1967; Lacey Jr. et al., 1992; Fitch and Upper, 1987; Higgs, 2009) if taken only in themselves.

The cost of this work was fully borne by the authors.

The authors declare no conflict of interest.

